# Variant Evolution Graph: Can We Infer How SARS-CoV-2 Variants are Evolving?

**DOI:** 10.1101/2024.09.13.612805

**Authors:** Badhan Das, Lenwood S. Heath

**Affiliations:** Department of Computer Science, Virginia Polytechnic Institute and State University, Blacksburg, Virginia, USA

**Keywords:** SARS-CoV-2, variant, evolution, graph, mutational distance, GISAID, virus

## Abstract

The SARS-CoV-2 virus has undergone mutations over time, leading to genetic diversity among circulating viral strains. This genetic diversity can affect the characteristics of the virus, including its transmissibility and the severity of symptoms in infected individuals. During the pandemic, this frequent mutation creates an enormous cloud of variants known as viral quasispecies. Most variation is lost due to the tight bottlenecks imposed by transmission and survival. Advancements in next-generation sequencing have facilitated the rapid and cost-effective production of complete viral genomes, enabling the ongoing monitoring of the evolution of the SARS-CoV-2 genome. However, inferring a reliable phylogeny from GISAID (the Global Initiative on Sharing All Influenza Data) is daunting due to the vast number of sequences. In the face of this complexity, this research proposes a new method of representing the evolutionary and epidemiological relationships among the SARS-CoV-2 variants inspired by quasispecies theory. We aim to build a Variant Evolution Graph (VEG), a novel way to model viral evolution in a local pandemic region based on the mutational distance of the genotypes of the variants. VEG is a directed acyclic graph and not necessarily a tree because a variant can evolve from more than one variant; here, the vertices represent the genotypes of the variants associated with their human hosts, and the edges represent the evolutionary relationships among these variants. A disease transmission network, DTN, which represents the transmission relationships among the hosts, is also proposed and derived from the VEG. We downloaded the genotypes of the variants recorded in GISAID, which are complete, have high coverage, and have a complete collection date from five countries: Somalia (22), Bhutan (102), Hungary (581), Iran (1334), and Nepal (1719). We ran our algorithm on these datasets to get the evolution history of the variants, build the variant evolution graph represented by the adjacency matrix, and infer the disease transmission network. Our research represents a novel and unprecedented contribution to the field of viral evolution, offering new insights and approaches not explored in prior studies.

## Introduction

The COVID-19 pandemic is caused by severe acute respiratory syndrome coronavirus 2 (SARS-CoV-2), a beta-coronavirus similar to the human SARS-CoV virus that caused the SARS outbreak (2002-2004) [1]. SARS-CoV-2 has caused millions of deaths and substantial morbidity worldwide. The pandemic generated unprecedented genetic data for a single pathogen [2], aiding in combating and understanding the virus’s biology. We saw evolutionary events previously limited to indirect inference, such as the diversification of SARS-CoV-2 into variants with myriad phenotypic traits like transmissibility, severity, and immune evasion [1].

Like other RNA viruses, coronaviruses evolve quickly, typically over months or years, with obvious and quantitative results. Evolution occurs in time frames equivalent to viral transmission events and ecological processes. As a result, evolutionary, ecological, and epidemiological processes impact each other in the study of RNA viruses [3]. Viral evolution is influenced by the pace at which mutations are generated and transmitted with populations. Natural selection will fix favorable mutations, such as the D614G mutation, which increases transmissibility [4]. Viral evolution adds another layer of complexity since viruses multiply and evolve within individuals while effectively transmitting from person to person, resulting in evolution on a different scale. Most variation is lost during the tight bottlenecks imposed at transmission, but other changes are often passed on by chance without selective benefit [5]. In addition to population-level dynamics, when viral lineages evolve, possibly resulting in antigenically different strains, higher-level processes such as lineage competition and extinction arise [1].

A phylogenetic tree or network is commonly used to illustrate the evolutionary history of a group of species, and this model has immensely aided in hypothesis development and testing. The SARS-CoV-2 genomic data set constantly grows as additional genomes are sequenced [6]. This expansion of the data set means that the phylogeny of SARS-CoV-2 has to be regularly updated [7], and the size of these data poses significant computational challenges for complete phylogenetic analysis [6]. According to Morel et al., [8], it is difficult to infer a reliable phylogeny on the GISAID data due to the enormous number of sequences and the small number of mutations. Furthermore, rooting the predicted phylogeny with some confidence, either through the bat and pangolin out-groups or by applying fresh computational methods to the in-group phylogeny, does not appear to be plausible [8]. Additionally, employing different multiple sequence alignment (MSA) strategies impacts the result of downstream phylogenetic analyses [9–11].

Viral quasispecies, also known as mutant spectra, swarms, or clouds, occur during the reproduction of RNA viruses in infected hosts [12, 13]. The concept of quasispecies stems from a speculative molecular evolution model that highlights the importance of error-prone, complex, and dynamic replication of primary RNA or RNA-like replicons in early life forms’ self-organization and flexibility [14, 15]. Quasispecies theory outlines the development of an effectively infinite population of asexual replicators with high mutation rates [16]. Due to that, the commonly utilized model for understanding viral evolution in a host is the theory of quasispecies [17, 18].

The overall picture of viral evolution somewhat differs from a phylogenetic tree or network because the ancestors from which new variants are evolving coexist in the same population, creating a cloud of variants. So, instead of using the phylogenetic tree or network to represent viral evolution, we introduce a graph data structure inspired by the quasispecies theory. It is an accepted model for viral evolution [12, 18–25]. Quasispecies is a continuous dynamic model closely associated with population dynamics [26]. A viral genotype is an RNA sequence. The set of all possible genotypes constitutes a genome sequence space of exponential size. All possible genotypes that we find theoretically cannot exist in the real world due to the bottlenecks of transmission and survival. The ones that survive through these bottlenecks are found in infected hosts. We may download the genotypes of these strains from a sequence database such as GISAID.

This paper proposes a Variant Evolution Graph, VEG, a novel way to model viral evolution. The core idea of this work lies in the quasispecies theory, which distinguishes it from existing literature and establishes a new direction for future research. In VEG, the vertices are the genotypes of the variants with associated hosts, and the edges represent the mutational distance and direction. We model genotype evolution as paths in this graph. During a pandemic, it is vital to have a good overview of the whole situation, and tracking the evolution of this pathogen in real-time can help understand its mechanisms, anticipate future trajectories, and develop preventative and treatment strategies [1]. VEG provides an overall picture of the evolution of the virus and how the variants are related to each other based on mutation. We also propose a disease transmission network, DTN, derived from this VEG, which infers the transmission directions among the hosts of a specific location, picturing the epidemiological scenario. Thus, our model gives the whole picture of both evolution and epidemiology. Due to the isolation protocol, we downloaded genome sequences from GISAID country-wise; Somalia (22), Bhutan (102), Hungary (581), Iran (1334), and Nepal (1719) are the five countries we used to download the genomes and run our proposed algorithm to get both VEG and DTN.

## Preliminaries

### SARS-CoV-2 Viral Genome

The development of SARS-CoV-2 has resulted in a terrible pandemic [27]. Coronaviruses are single-stranded, enveloped RNA viruses [28]. The genome sequence is 29, 903 bp long. The RNA genome contains several genes that encode various proteins essential for the virus’s replication, transcription, and infection processes. Fifteen key genes found in the SARS-CoV-2 genome according to GISAID are shown in Table 1.

**Table 1.** Fifteen key genes of the SARS-CoV-2 genome and their locations in the reference genome of SARS-CoV-2 isolate Wuhan-Hu-1 [29].

| Gene | Location |
| --- | --- |
| ORF1ab [29–31] | 266..21555 |
| ORF1a [29–31] | 266..13468 |
| S [30, 31] | 21563..25384 |
| ORF3a [30, 31] | 25393..26220 |
| ORF3b [30, 31] | 25765..26220 |
| E | 26245..26472 |
| M [30, 31] | 26523..27191 |
| ORF6 [30, 31] | 27202..27387 |
| ORF7a [30, 31] | 27394..27759 |
| ORF7b [30, 31] | 27756..27887 |
| ORF8 [30, 31] | 27894..28259 |
| N [30, 31] | 28274..29533 |
| ORF9b [30, 31] | 28284..28577 |
| ORF9c [30] | 28734..28955 |
| ORF10 | 29558..29674 |

### Average Nucleotide Identity

Average nucleotide identity (ANI) is a measure of nucleotide-level genomic similarity between the coding regions of two genomes [32]. Goris et al. (2007) introduced the generally recognized notion of ANI as an alternative to DDH (DNA-DNA Hybridization) by emulating the DDH experimental approach [33]. It yields a value between 0 and 1, generally expressed as a percentage. Typically, an ANI threshold of 95 *−* 96% corresponds well with the 70% DDH value historically used to define species boundaries, making ANI a frequently used standard for microbial taxonomy [34].

### Sourmash

Sourmash is a software tool for comparing and analyzing extensive collections of genomes or other biological sequences, such as transcriptomes or metagenomes [35, 36]. It compares sequences based on *k*-mer content, which are short, fixed-length substrings of sequences. MinHash is a technique for generating signatures and compact representations of *k*-mer contents in sourmash. These signatures are generated using hashing techniques and contain information about the presence and abundance of specific *k*-mer within the sequences. Fractional MinHash or FracMinHash is a refined technique used to create more compact and efficient MinHash sketches while maintaining the accuracy of similarity estimates. Sourmash (v4.4) can estimate ANI between two FracMinHash (scaled) sketches [37].

### Pyani

Pyani is a program and Python package that supports calculating ANI and related measures for whole genome comparisons and providing relevant graphical and tabular summary outputs [38]. Pyani provides four sub-commands to run ANI analyses: anim (ANIm, using MUMmer software package for sequence alignment), anib (ANIb, using BLAST+), aniblastall (ANIb, using legacy BLAST), and tetra (using tetranucleotide frequency correlation coefficients as a measure of genomic similarity). We have used anib for our ANI calculation.

### Levenshtein (Edit) Distance

Levenshtein distance [39], also known as edit distance, is a mathematical distance or metric between two strings. The edit distance between two strings is measured by the minimum cost sequence of edit operations needed to change one string into another. The edit operations include changing one symbol of a string into another single symbol (substitution), deleting one symbol from a string (deletion), or inserting a single symbol into a string (insertion). Edit distance provides a quantitative means to compare and align sequences.

### Defining Mutational Distance

We will define two types of values as mutational distance: (1-ANI) and edit distance. In our methods, we call both terms mutational distance for ease of discussion. Although the edit distance of two genome sequences logically represents the mutational distance between those two genotypes, ANI is a similarity measure. Since we need mutational distance, we use the value (1-ANI), which can be perceived as the measure of dissimilarity between two genomes. (1-ANI) is not a mathematical distance, but we will use the term ‘mutational distance’ for both (1-ANI) and edit distance for convenience.

### Introducing Variant Evolution Graph

In a specific location, let us consider a virus mutating and, thus, evolving. In a certain span of time, *n* variants have evolved, where each of these variants is associated with an evolution (collection) date. A variant evolution graph, *V EG* = (*V, E, w*), is a directed acyclic graph, where vertex set, *V* represents the *n* strains, edge set, *E* = {(*u, v*)|v strain evolved from u*}* is the set of weighted edges signifying the evolution directions and portraying the parent-child relationships, and weight function, *w* : *E →* ℝ represents the mutational distance.

### Problem Statement

Let *L* be a genome space of the genotypes of *n* strains of that virus in a specific location associated with their collection dates.

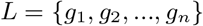

Here, *g*_*i*_ is the *i*^*th*^ genome in set *L*, where collection dates sort the genomes.

We have a *n × n* distance matrix, *D*, where *D*_*ij*_ is the mutational distance between the *i*^*th*^ and *j*^*th*^ genomes; 1 *≤ i, j ≤ n*. Our first objective is to build the Variant Evolution Graph, *V EG* = (*V, E, w*) based on the mutational distances among these strains. Our second objective is to derive the disease transmission network, DTN, a subgraph of VEG, portraying the direct transmissions among the human hosts of that specific location.

## Materials and Methods

### Data Set

For building VEG, we have used the genotypes of the SARS-CoV-2 variants. We downloaded the complete reference genome of SARS-CoV-2 isolate Wuhan-Hu-1 from NCBI [29] and the genome sequences of the strains from GISAID that are complete, high-coverage and have a collection date. The genome space for building the graph was decided country-wise or state-wise, depending on the location due to the isolation protocol during the COVID-19 pandemic. As mentioned, we downloaded variant genomes from Somalia, Bhutan, Hungary, Iran, and Nepal. In our method, we kept a variable, *τ*, that represents the cumulative threshold of the percentage of Ns in the genomes of a particular data set. We have used *τ* = 0 for our experiments, so we have only chosen genomes with no Ns in their sequences. Table 2 shows the count of genomes before and after this filtering.

**Table 2.**
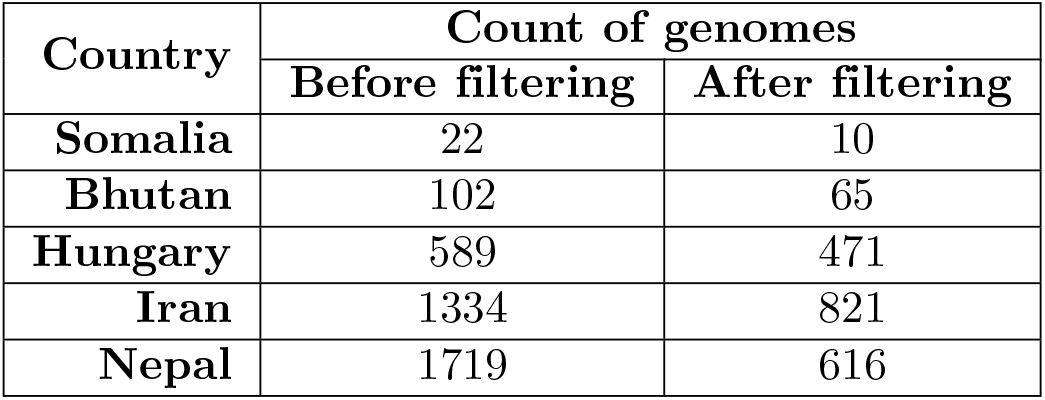
Table showing the counts of genomes before and after filtering based on *τ* = 0 for Somalia, Bhutan, Hungary, Iran, and Nepal data sets.

| Country | Count of genomes |  |
| --- | --- | --- |
|  | Before filtering | After filtering |
| Somalia | 22 | 10 |
| Bhutan | 102 | 65 |
| Hungary | 589 | 471 |
| Iran | 1334 | 821 |
| Nepal | 1719 | 616 |

### Edit Distance Calculation

According to the positions of the genes, we considered ORF1ab, S, ORF3a, E, M, ORF6, ORF7a, ORF7b, ORF8, N, and ORF10 to generate the coding region for the calculation of edit distance. The locations of these genes in the reference genome are given in Table. 1. As shown in Fig 1, initially, we take all the key genes and then run BLASTn against the *n* variant genomes. This results in eleven extracted gene sequences per genome, which later are concatenated sequentially to be considered as coding sequences.

**Fig 1.**
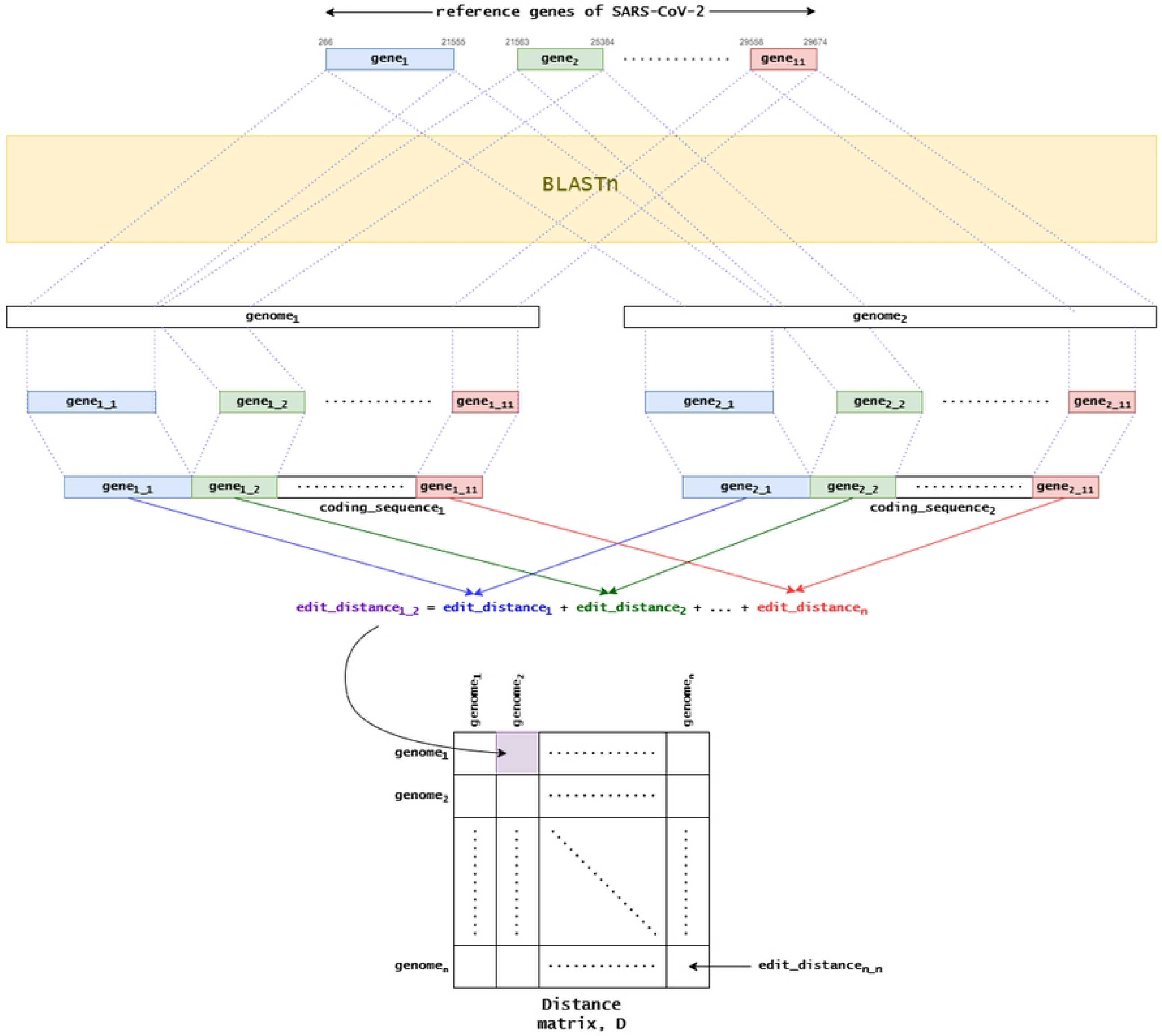
Pipeline for generating distance matrix using edit distances between the variant genomes. Here, *gene*_*i j*_ represents the *j*^*th*^ gene in the *i*^*th*^ genome; 1 *≤ i ≤ n* and 1 *≤ j ≤* 11. The element at row *i* and column *j, D*_*ij*_, is the edit distance between *genome*_*i*_ and *genome*_*j*_, for 1 *≤ i, j ≤ n*.

### Algorithms

#### Assumptions

For our method, we have some assumptions to start with. Those are as follows.

1. The collection date associated with a specific variant is assumed to be when that variant first appeared in the population.
2. Multiple variants having the same collection date cannot evolve from each other.

#### Computing Distance Matrix

We ran Sourmash and Pyani on the genome sequences of a genome space to get the distance matrices: *D*_*S*_ and *D*_*P*_. Sourmash gives a distance matrix where the elements hold the (1-ANI) values, and we use that matrix as *D*_*S*_. Pyani returns an ANI percentage identity matrix, *D*_*I*_, where the elements hold the ANI values. Since we use the dissimilarity value (1-ANI) as mutational distance, we then process *D*_*I*_ to a distance matrix, *D*_*P*_, where *D*_*I*_ = 1 *− D*_*P*_.

We focused on the coding region rather than the whole genome sequence to compute the edit distance for the distance matrix. As shown in Fig 1, for two genomes, *genome*_1_ and *genome*_2_, edit distances are computed at the gene sequence level and then added to calculate the total edit distance of these two genomes. We compute all the edit distances between pairwise genomes and generate the edit distance matrix, *D*_*E*_.

#### Building the Variant Evolution Graph

The genome set, *L*, is sorted in ascending order according to the collection dates of the genomes from the oldest to the newest variants. As a result, the distance matrix, *D*, is also sorted according to their collection dates. According to the second assumption in Section, *D* is organized using the Algorithm 1. This ensures we do not compare the distances of two genomes collected on the same date.

##### Algorithm 1 Organize-Distance-Matrix(*D, L, dates*)

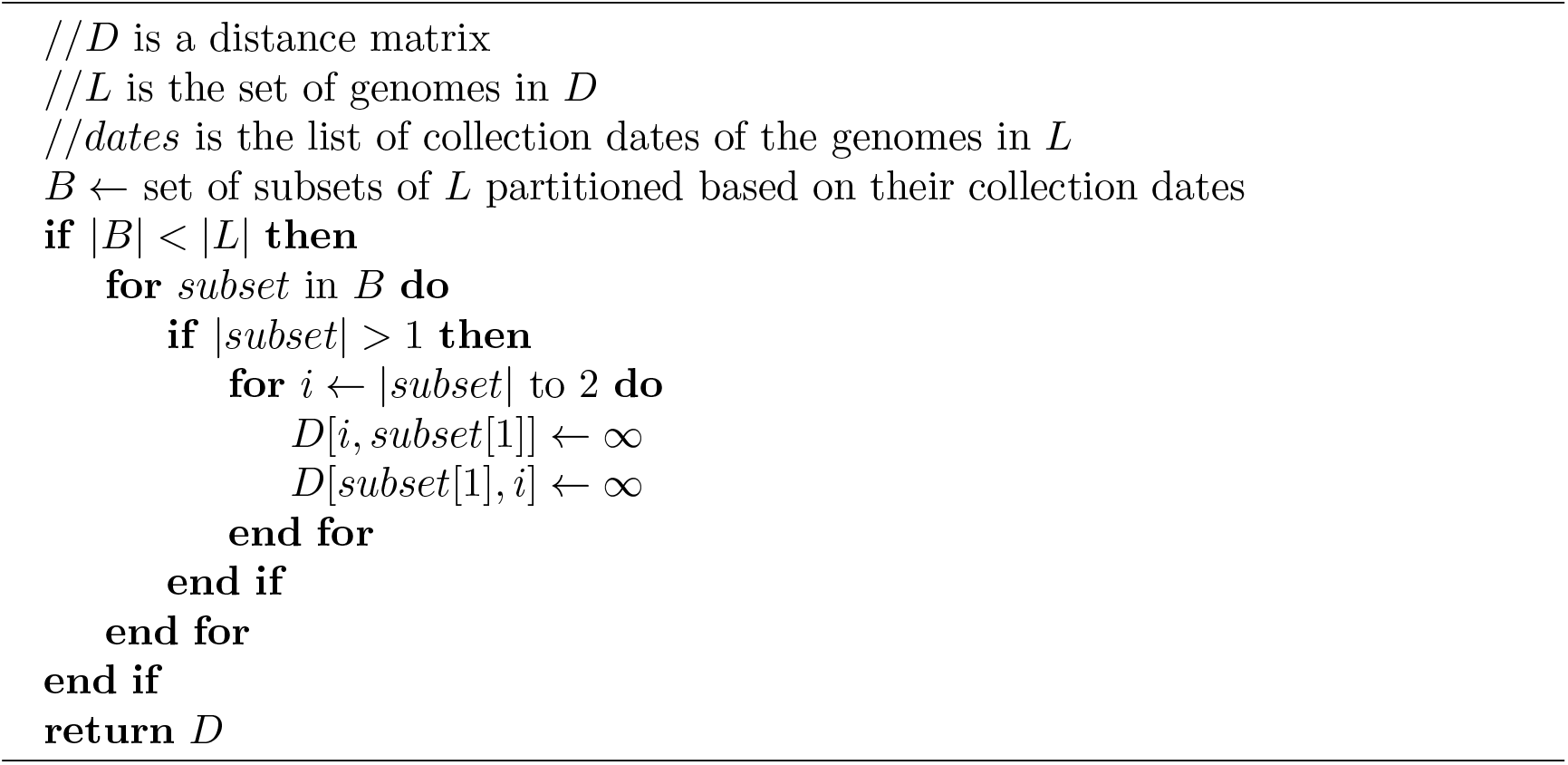

Suppose for *n* variant genomes, there are *B* different collection dates, where *B ≤ n*. So, we have *B* partitions of genomes grouped by their collection dates. Since these *n* genomes are sorted in ascending order of their collection dates in *L*, the *B* groups are also sorted accordingly. The variants of the *i*^*th*^ group can only evolve from the variants of the groups having earlier collection dates, where 1 *< i ≤ d*. For the first group, where *i* = 1, the variant(s) are the oldest, and those do not have any parents. As shown in Algorithm 1, all the pairwise distances between the strains of a group are set to *∞*. This is done for the second assumption, that the variants with the same collection date cannot evolve from each other.

As shown in Algorithm 2, the main idea of this method is to trace back the evolutionary pathways from the recent variants to the oldest ones by detecting the closest variant(s) from each variant. The closest variant(s) of a variant *v* has the smallest mutational distance from *v*, and we infer that this closest-distance variant(s) is the one from which *v* has evolved. Let us consider *P* as the set of variants from which *v* has the smallest distance. Reversing this relationship, we can call *v* as the child of each variant in *P*. We get these child-parent relationships for all the variants existing. These relationships are then converted to the edges of a graph, *G* = (*V, E, w*), where *V* = *L*. In the edge set, *E*, the weight, *w*, of an edge (*u, v*) is the mutational distance between the variants *u* and *v*. So, the weight function, *w* = {*D*_*uv*_ |(*u, v*) *∈ E*}. These relationships could have formed a tree structure, but some variants might have more than one parent, creating a directed acyclic graph (DAG) but not necessarily a tree. All the evolutionary pathways are modeled as paths in this graph. The entire pipeline of generating VEG is shown in Fig 2. The Build-VEG method generates three graphs, *G*_*S*_ = (*V*_*S*_, *E*_*S*_, *w*_*S*_), *G*_*P*_ = (*V*_*P*_, *E*_*P*_, *w*_*P*_), and *G*_*E*_ = (*V*_*E*_, *E*_*E*_, *w*_*E*_) for the three methods, sourmash, pyani, and edit distance, respectively.

**Fig 2.**
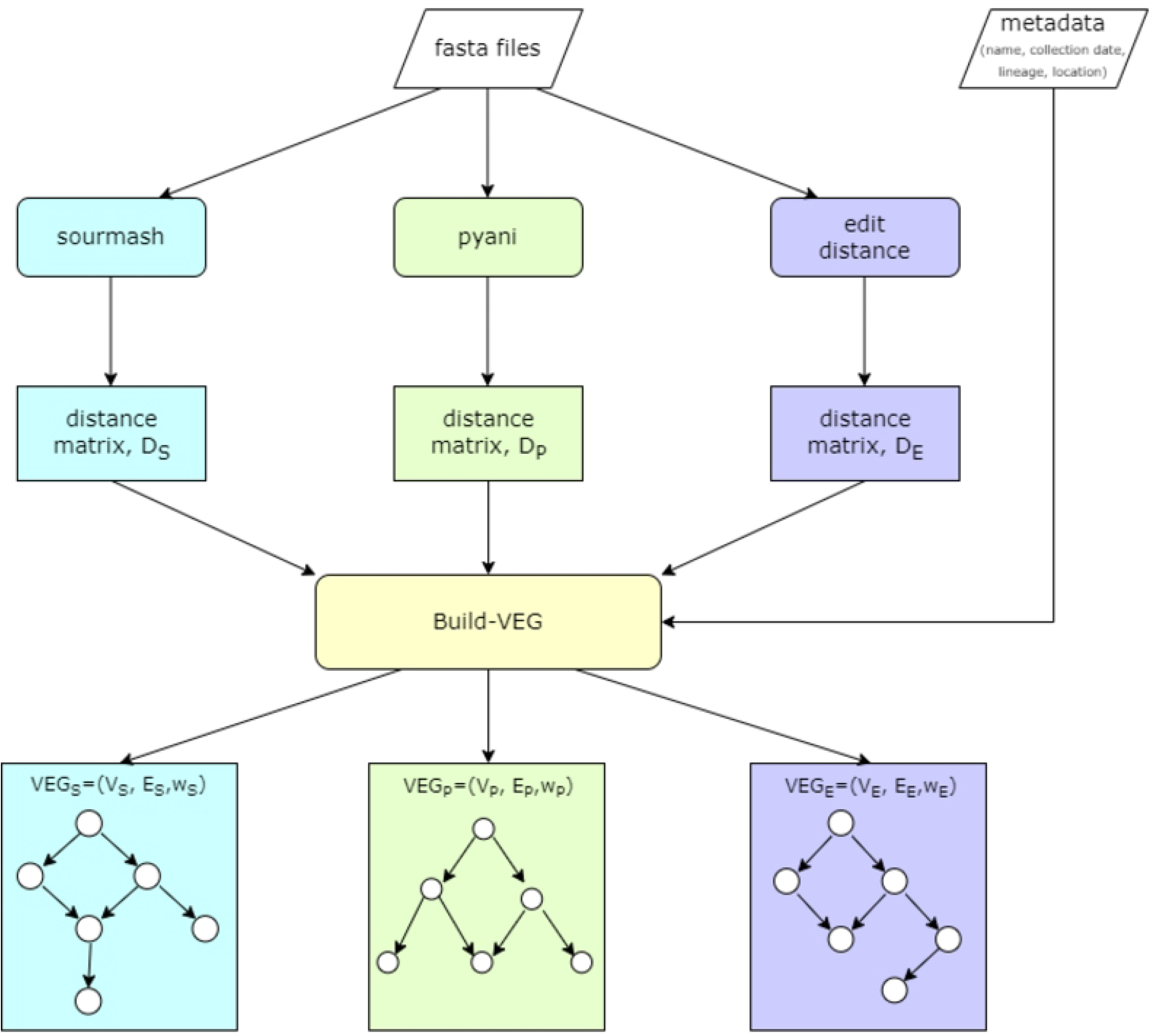
The whole pipeline of building VEG. The edit distance computation in this pipeline is separately shown in Fig 1.

##### Algorithm 2 Build-VEG(*D, L*)

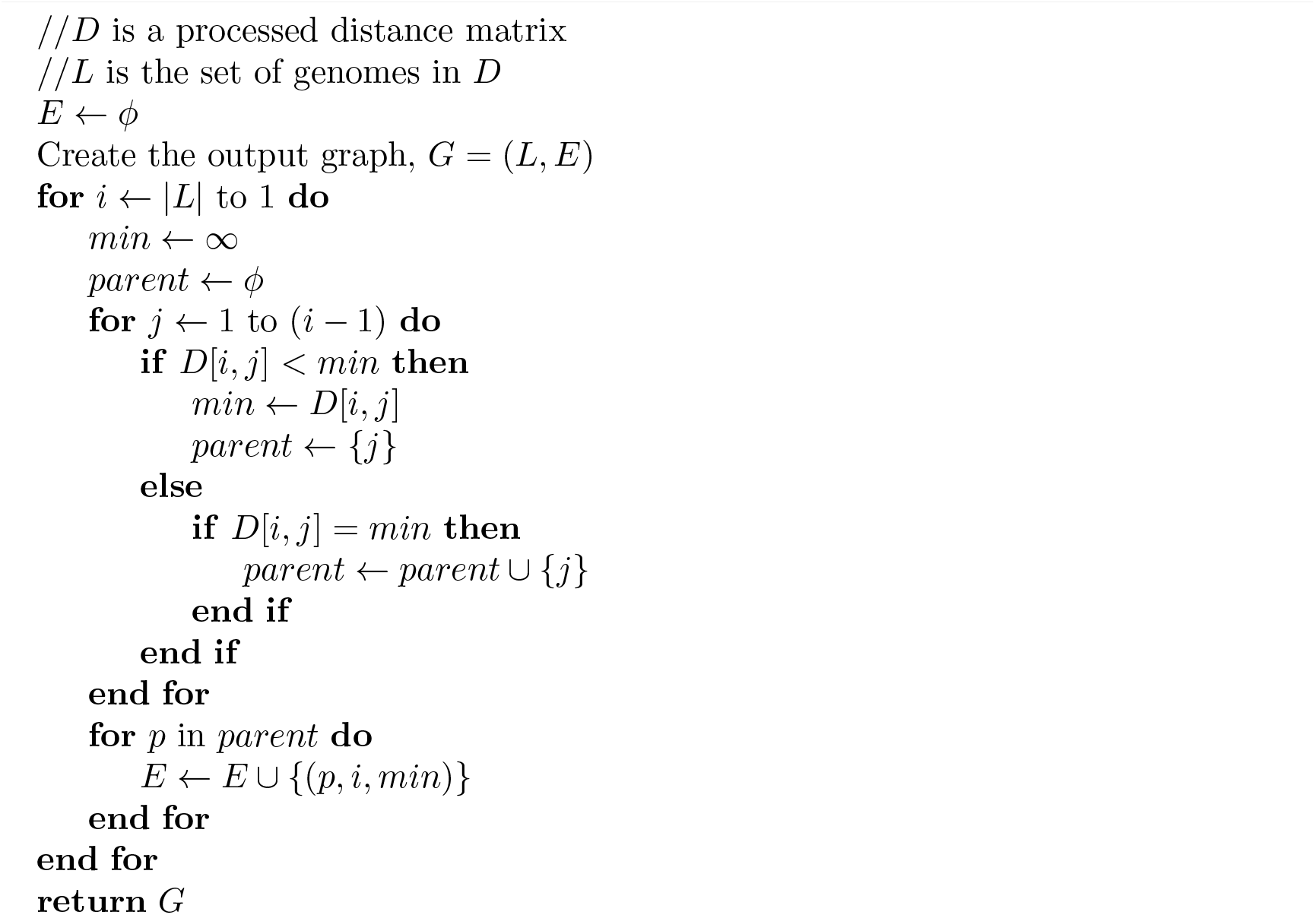

### An Example

Let us consider a small example of a genome space with six viral genomes: *A, B, C, D, E*, and *F*, shown in Fig 3. Each of these genomes is associated with a collection date. According to assumption 1 in Section, the collection dates are assumed to be the dates when these variants first appeared in the population. We sort the rows and columns of the distance matrix (from sourmash, pyani, or edit distance) in the ascending order of the collection dates. In this example, variants *E* and *F* have the same collection date, June 7, 2020. So, according to assumption 2 in Section, *E* and *F* cannot evolve from each other. So, we set the distance at position *D*_*FE*_ = *∞* and *D*_*EF*_ = *∞*. In this example, we are using the edit distance matrix.

**Fig 3.**
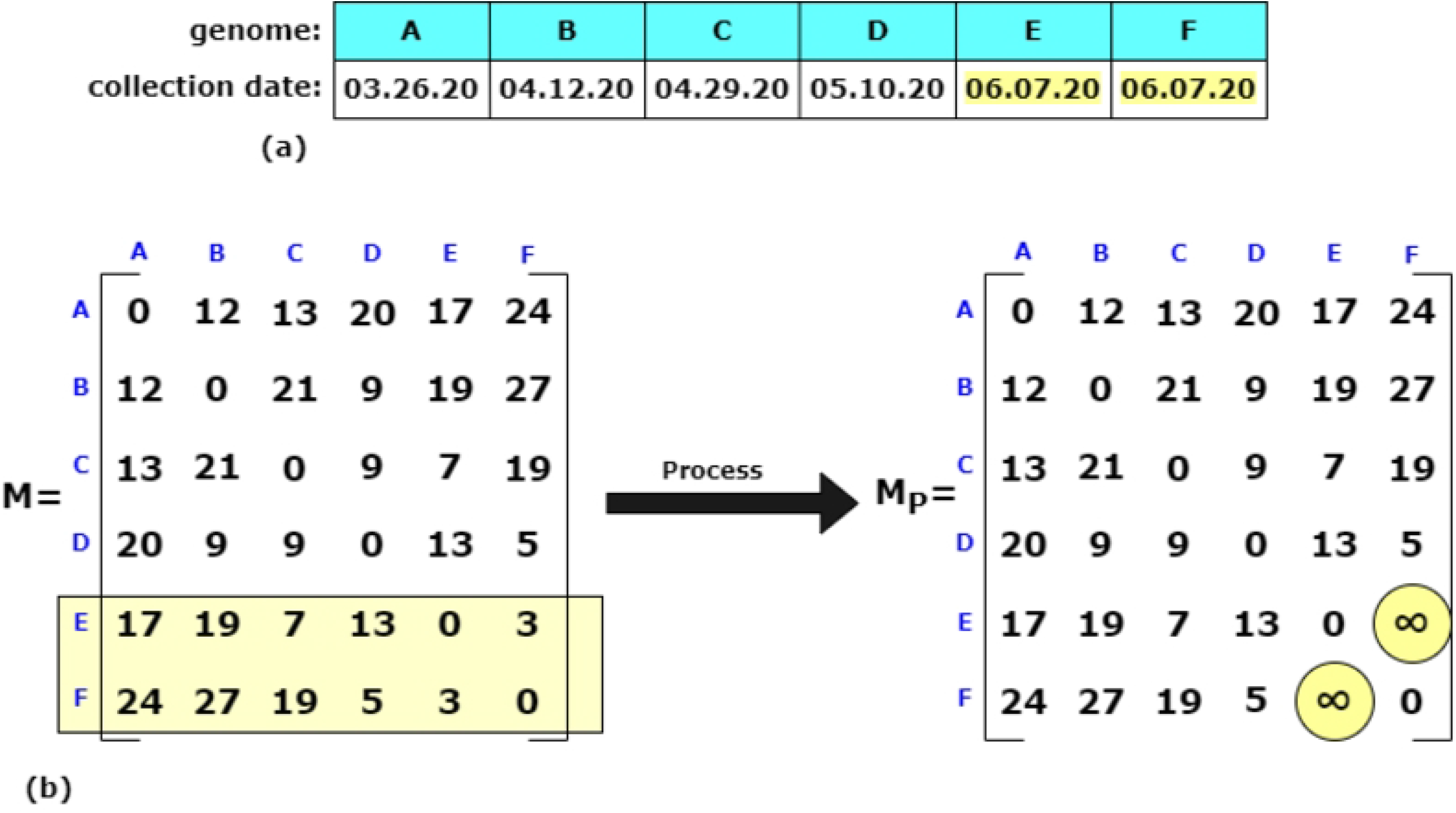
A sample set of six genomes. (a) The set of six variant genomes, *A, B, C, D, E*, and *F*, and their corresponding collection dates. (b) The workflow of Algorithm 1 on the distance matrix, *M*.

Next, we pass this distance matrix to Build-VEG algorithm along with the set of genomes, *L* = {*A, B, C, D, E, F*}. The smallest distance(s) in each row is marked with blue circles, shown in Fig 4. Some variants might have more than one cell with the smallest distance; for example, for *D*, we find that 9 is the smallest distance, and *D*_*DB*_ = *D*_*DC*_ = 9. So, according to our method, genome *D* has two parents: *B* and *C*. Identifying the smallest distances from each strain, we filter out the relationships and list the parent sets for each. Let *G*_*E*_ = (*V*_*E*_, *E*_*E*_, *w*_*E*_) be our VEG, where *V*_*E*_ = *L*. Since we are tracing the parent(s) from each child strain, we reverse these relationships to get the (parent, child) relationships for the edge set, *E*_*E*_. Here, *w*_*E*_ = {*D*_*uv*_ |(*u, v*) *∈ E*_*E*_}.

**Fig 4.**
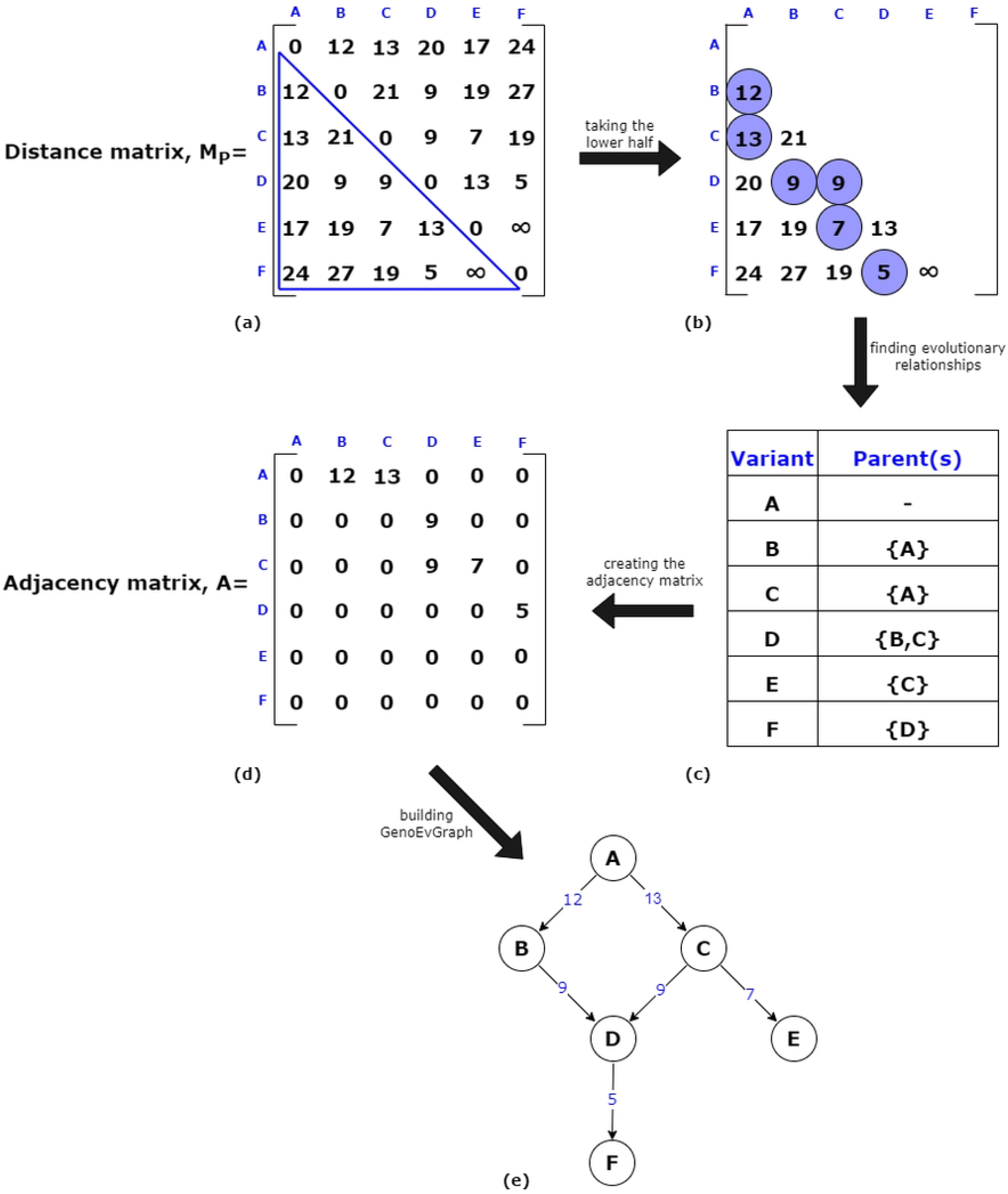
The workflow of Algorithm 2 with the example in Fig 3.

### Deriving the Distance Transmission Network

Since VEG is constructed based on mutation and evolution timeline, it provides the evolutionary relationships among the strains in a genome space as well as infers the DTN, of that specific location since each of these strains is associated with a patient or host. The edges of the VEG also show day differences between the collection dates of the strain genomes since each node has a collection date. The infectious period is between seven and ten days [40, 41]; edges with day differences of less than eleven days can be considered direct transmissions between the hosts. For an edge (*u, v*), let us consider the collection dates of *u* and *v*, which are *d*_1_ and *d*_2_ respectively. Then, Δ*d* = |*d*_2_ *− d*_1_| is the day difference of the edge (*u, v*). So, any edge having Δ*d ≤* 10 can be considered as direct transmission between the two hosts of that edge.

## Results

### Count of N’s

As mentioned earlier in Section, the count of N’s, unknown nucleotides, is an important parameter to be considered while processing the data set. After several experiments, we found that the complete, high-coverage genomes in GISAID have at most 1% Ns in their sequences. Fig 5 shows the change in the count of filtered genomes by changing the threshold, *τ*, for Somalia, Bhutan, Iran, and Nepal data sets, respectively. However, all our experiments are done setting *τ* = 0, which means we are only considering those genomes that do not have any Ns in their sequences.

**Fig 5.**
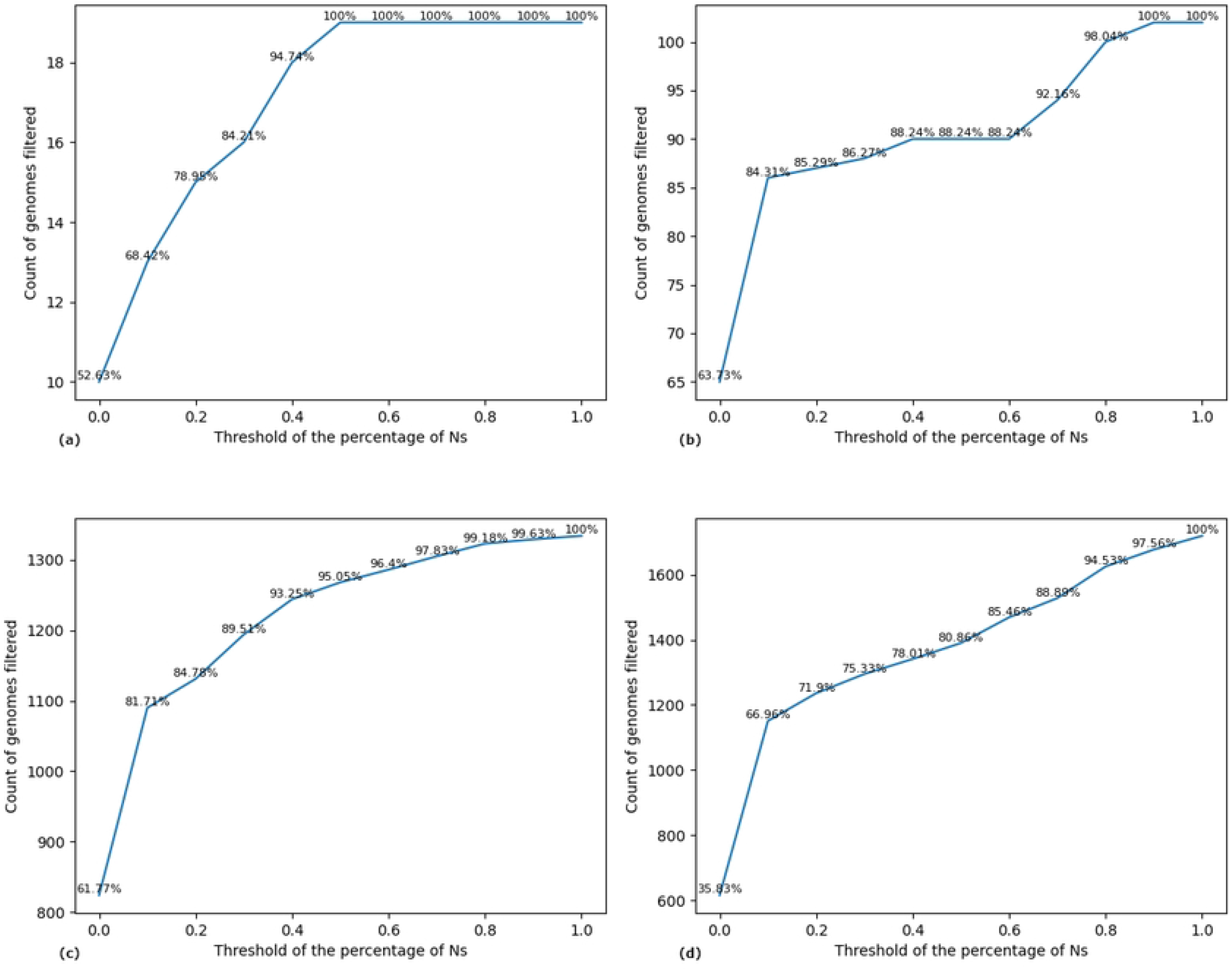
Count of genomes filtered based on the percentage of Ns, *τ*, in the genomes. The x-axis shows the threshold values, and the y-axis shows the count of filtered genomes. The plots are of (a) Somalia, (b) Bhutan, (c) Iran, and (d) Nepal data sets.

We found a significant difference between the count of Ns in the whole genome sequence and the coding region. Although we have chosen *τ* = 0 to build our graph, *τ* can be any other value. For *τ* = 0, there are no Ns in the genome sequences and, thus, no Ns in their corresponding coding regions. But, in general, coding regions have fewer Ns than whole genomes. So, working with coding regions gives more accurate results than with entire genomes from the perspective of the existing Ns in the sequences. Fig 6 shows the average count of Ns across the genome as the whole sequences vs the coding regions in Somalia, Bhutan, Hungary, Iran, and Nepal data sets. It is apparent from Fig 6 that the coding region has significantly less Ns count than the whole genome.

**Fig 6.**
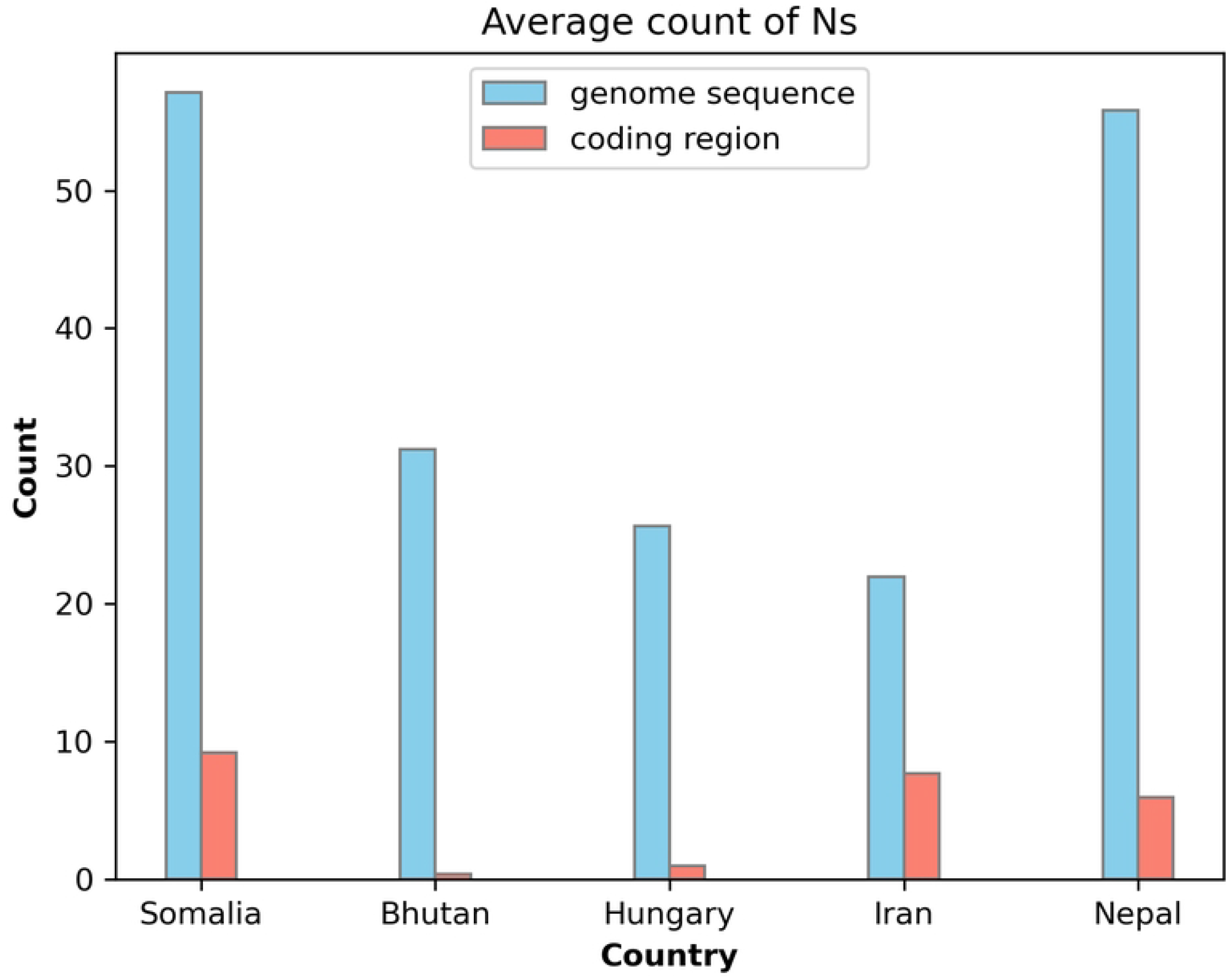
Average count of Ns in the genome sequences vs the coding regions in five data sets.

### Limitations in Data Sets

During the pandemic, only some of the infected people tested themselves, and as a result, GISAID could not gather a good number of patient data. So, the edges that have a day difference, Δ*d >* 10, can be viewed as a chain of transmissions. Moreover, to compute the mutational distance as accurately as possible, we only downloaded strains with complete, high-coverage genomes with complete collection dates from GISAID. This downloaded set was further filtered based on the count of Ns, where we kept the ones with no Ns. So, a bigger portion of the data set has been filtered out to be precise about the mutational distance.

### Variant Evolution Graph

We aim to build a directed and weighted VEG based on the mutational distance. We have used sourmash, pyani, and edit distance to compute the distance matrices. As a result, our method produces three graphs. We have used the adjacency matrix for the graph representation. For each type mentioned, our algorithm gives the following outputs. The VEG for the Bhutan data set using the edit distance matrix is shown in Fig 7.

**Fig 7.**
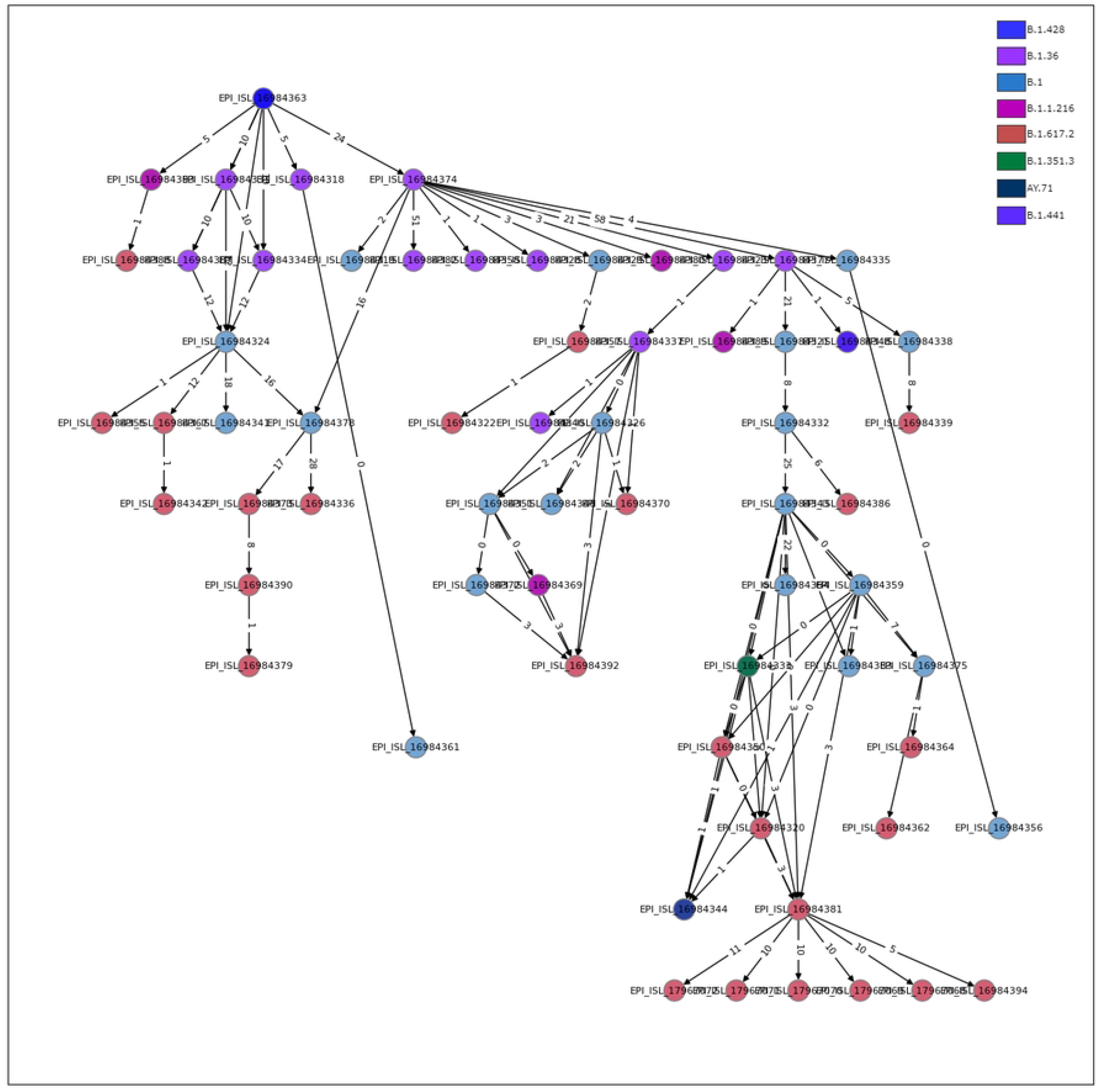
VEG of Bhutan data set using the edit distance matrix. The color code for lineages is shown in the top-right corner.

- **Evolution history log**: The evolution history log has the following information for each variant in the data set.
  - set of parent variants
  - mutational distances from the parent variants
  - day differences from the collection dates of the parent variants
- **Adjacency matrix**
- **Variant Evolution graph**: a.png file; variants of the same lineage have the same color (colors chosen randomly)
- **Lineage information**: All the variants are grouped by their PANGO [42] lineages (provided by GISAID).

### Comparison Among Sourmash, Pyani, and Edit Distance

As described in Section, we have built VEG based on mutational distance derived from sourmash, pyani, and edit distance. So, for each country data set, we get three VEGss: *G*_*S*_ = (*V*_*S*_, *E*_*S*_, *w*_*S*_), *G*_*P*_ = (*V*_*P*_, *E*_*P*_, *w*_*P*_), and *G*_*E*_ = (*V*_*E*_, *E*_*E*_, *w*_*E*_), as shown in Fig 2. We compared these three graphs by finding out how many parent-child mutational relationships they have in common. This can easily be tracked down by the edges they have in common. However, this will not give an overall picture of the evolutionary pathways.

The Venn diagrams in Fig 8 show how similar the parent-child relationships are in Bhutan, Hungary, Nepal, and Iran data sets. From these Venn diagrams, we can perceive that sourmash, pyani, and edit distance are inferring different parent-child relationships. Even though sourmash and pyani estimate ANI, their inferred relationships are pretty different, as seen in these results. Edit distance calculation is more reliable and accurate for the mutational distance since we get the nucleotide difference between two sequences. Sourmash and pyani do not agree regarding ANI values among the variants’ genomes. In contrast, edit distance becomes a promising measurement for mutational distance and, as a result, gives better results than both sourmash and pyani.

**Fig 8.**
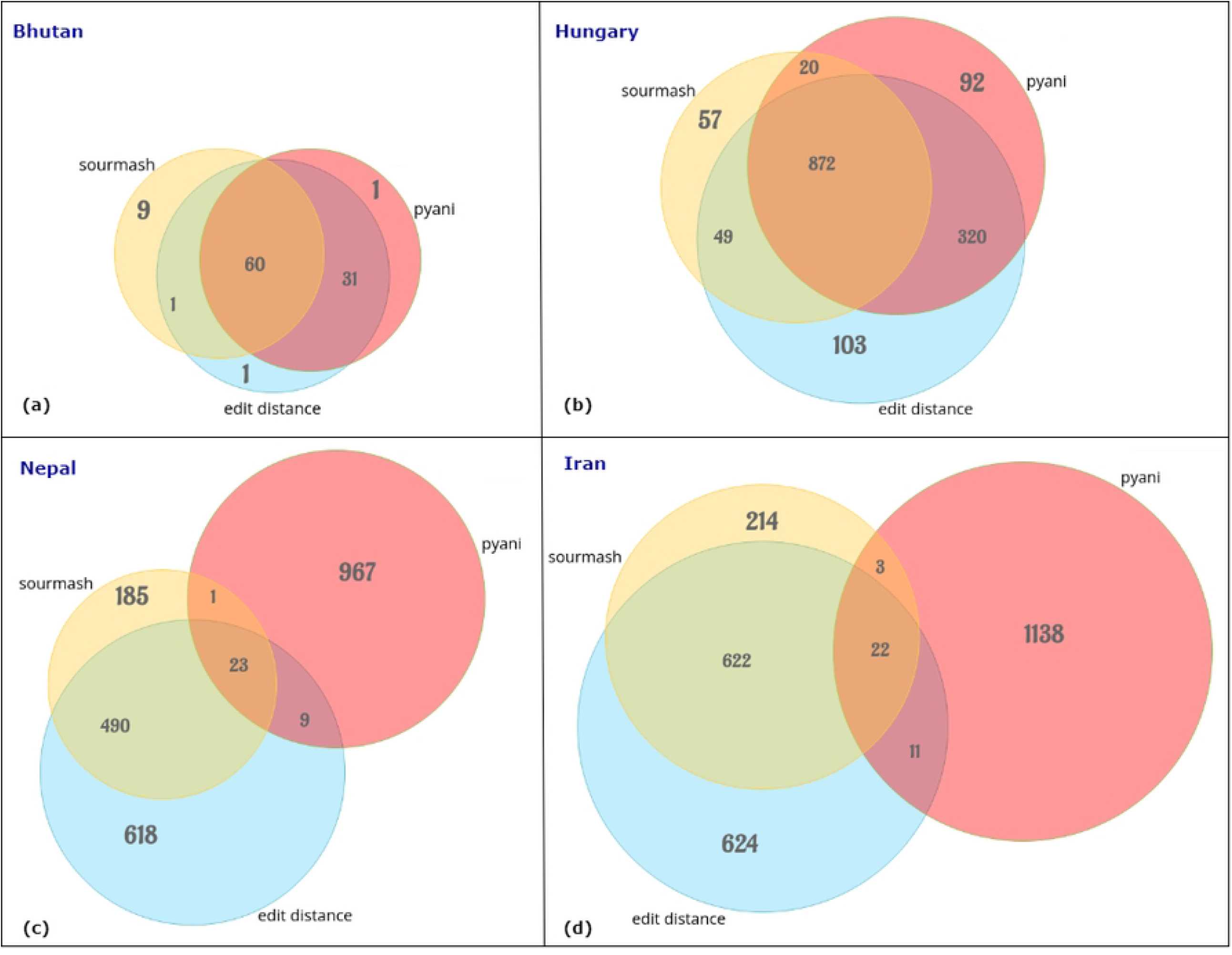
Venn diagrams showing parent-child relationships among the VEGs derived from sourmash, pyani, and edit distance. (a) Bhutan, (b) Hungary, (c) Nepal, and (d) Iran data sets.

### Disease Transmission Network

We can infer the DTN from the VEG described in Section. Fig 9 presents the inferred DTN of the Bhutan data set. It is important to note that DTN should ideally be connected during a pandemic if all the patients are tested on time and their variants are sequenced and stored in a database. But the DTN of Bhutan shown in Fig 9 is disconnected, which can be attributed to the real-world scenario of COVID-19. If a significant number of samples in GISAID exhibited the qualities mentioned in Section, the VEG and its inferred DTN would have been more comprehensive, with a greater number of variants, and a higher number of transmissions could have been inferred. The limitations mentioned in Section underscore the need for a more extensive and diverse data set for more accurate and comprehensive results.

**Fig 9.**
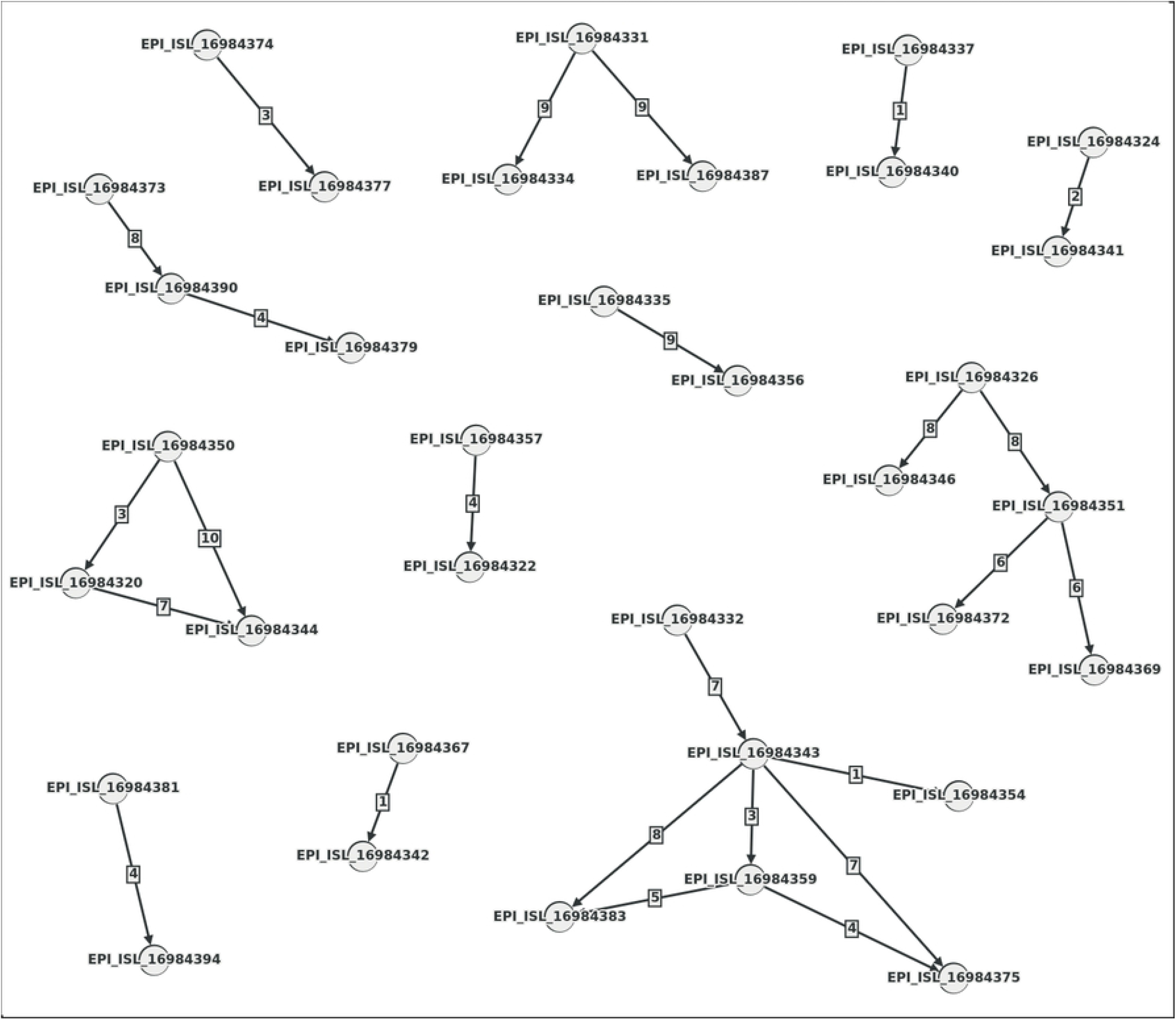
The DTN is inferred from the VEG of the Bhutan data set (edit distance). Here, the nodes are the hosts, and the edges represent the direction and day differences of the inferred transmissions.

### Identifying Superspreaders

Several studies have indicated that 20% of the host population has the potential to cause 80% of transmission occurrences, a pattern known as the 80*/*20 rule [43–47]. Understanding the role of super-spreaders might lead to more efficient disease outbreak containment and more accurate epidemic modeling [48].

In the DTN mentioned earlier, the nodes are the hosts, among which some superspreaders exist, which the out-degrees of each node can find. For a transmission network, we list the count of out-degrees of all the nodes and, sorting the list in descending order, we separate it into two parts: higher-degree set with the topmost 20% of the degrees and lower-degree set with the rest. According to the 80*/*20 rule, the nodes of this higher-degree set are responsible for 80% of the transmission, which means the nodes of this set have 80% of the total out-degrees.

Table 3 shows the statistics of Bhutan, Hungary, Iran, and Nepal data sets where the 80^*th*^ percentile is 1.0, 2.0, 1.0, and 1.0, respectively. This means all the out-degrees higher than these values belong to the higher 20%. Partitioning the out-degree lists of these data sets into the higher-degree and lower-degree sets, we found out that for the Bhutan data set, 60% of the out-degrees belong to the higher-degree set, 80.37% for the Hungary data set, 90.71% for the Iran data set, and 78.33% for the Hungary data set, as shown in Table 4. Some percentages do not exactly follow the 80*/*20 rule, which can be explained by the limitations mentioned earlier.

**Table 3.** Statistics showing the degree distribution of the vertices in disease transmission networks of Bhutan, Hungary, Iran, and Nepal.

|  | Bhutan | Hungary | Iran | Nepal |
| --- | --- | --- | --- | --- |
| count | 34 | 318 | 624 | 326 |
| mean | 0.735294 | 1.361635 | 1.466346 | 0.990798 |
| std | 0.931237 | 2.573471 | 4.690637 | 2.140433 |
| min | 0.000000 | 0.000000 | 0.000000 | 0.000000 |
| 20% | 0.000000 | 0.000000 | 0.000000 | 0.000000 |
| 40% | 0.000000 | 0.000000 | 0.000000 | 0.000000 |
| 50% | 0.500000 | 0.000000 | 0.000000 | 0.000000 |
| 60% | 1.000000 | 1.000000 | 0.000000 | 1.000000 |
| 80% | 1.000000 | 2.000000 | 1.000000 | 1.000000 |
| max | 4.000000 | 17.000000 | 62.000000 | 17.000000 |

**Table 4.**
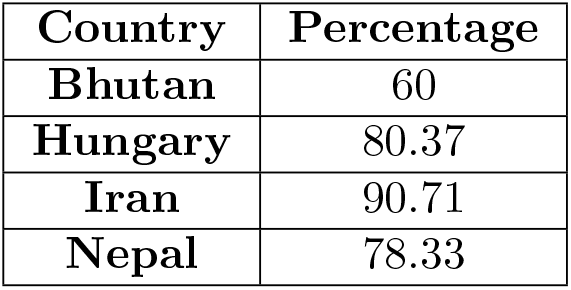
Percentages of the count of out-degrees in the higher-degree sets of Bhutan, Hungary, Iran, and Nepal data sets.

## Discussion and Conclusion

Our research is novel and unprecedented, filling a significant gap in the existing knowledge and advancing the field uniquely. We have primarily focused on the variants of SARS-CoV-2, a virus that has caused a severe pandemic and has a vast data set of variants. This virus has served as an excellent example of both evolution and epidemiology. However, the method we have developed is not limited to SARS-CoV-2. It is a versatile tool that can be applied to any viral outbreak, providing researchers and scientists with a comprehensive and early understanding of the situation and enabling them to take prompt and necessary actions.

We focused primarily on the genes for the edit distance calculation, so we only computed the edit distance for coding regions. However, this can also be done for the whole genome sequence. One way to do this is by creating blocks of 1000 nucleotides. Instead of genes, these blocks can be aligned against the reference genome, and the edit distance matrix can be computed similarly, as described in Fig 1. So, both ways, using edit distance to compute the mutational distance gives better and more accurate results than ANI.

Our method entirely depends on the quantity and quality of the genome sequences submitted to a global database for viral pandemics like GISAID. If the quality is not good enough, the quantity falls, creating discrepancies in the building of VEG and inferring the DTN. But, in any future pandemic, if samples are gathered and appropriately organized, our method can help visualize the variants’ evolution in evolutionary and epidemiological ways. For future direction, both VEG and DTN can be used to make machine learning models learn and predict potential variants and transmissions, thus helping researchers and medical professionals take necessary steps to prevent massive disastrous pandemic situations.

## Data Availability Statement

All SARS-CoV-2 genome sequences are publicly available from the GISAID database. The source code can be found at https://github.com/Badhan023/viral_evolution.

## Acknowledgments

We acknowledge the support of National Science Foundation grant number 1918656.

